# Optimizing efficacy to safety ratio of glucocorticoids in rheumatoid arthritis models by leveraging PPARα agonism

**DOI:** 10.1101/2025.10.14.682304

**Authors:** Lisa L. Koorneef, Elisabeth Gilis, Dorien Clarisse, Daria Fijalkowska, Sara Dufour, Julie Coudeneys, Annick Verhee, Jonathan Thommis, Matthias Kerkhofs, Guillaume Planckaert, Simon Devos, Leander Meuris, Dirk Elewaut, Karolien De Bosscher

## Abstract

**Objectives:** Glucocorticoids remain essential therapies for several immune- and inflammatory diseases such as rheumatoid arthritis (RA) but are notorious for their (metabolic) side effects. Given the anti-inflammatory and metabolically favorable actions of peroxisome-proliferator activated nuclear receptor (PPAR) agonists, we investigated whether PPARα agonism could enhance the therapeutic efficacy and/or mitigate the (metabolic) side effects of glucocorticoids.

**Methods:** We evaluated the effects of the synthetic glucocorticoid dexamethasone and the PPARα agonist GW7647 (GW) across three RA model systems: L929sA fibroblasts, primary human fibroblast-like synoviocytes (FLS) and collagen-induced arthritis (CIA) mice.

**Results:** Dexamethasone reduced the inflammatory TNFα response in L929sA cells, which was further potentiated by GW. In vivo however, GW reduced the dexamethasone-induced adiposity and hypertriglyceridemia, but not arthritis severity. Curiously, GW alone induced several proinflammatory genes within arthritic synovium which were counteracted by glucocorticoids. Proteomic profiling of TNFα-stimulated human FLS revealed that combined use of dexamethasone and GW selectively suppressed interferon-stimulated proteins. In line herewith, co-stimulation with TNFα and IFNβ amplified the suppressive effect of combined dexamethasone and GW treatment on pro-inflammatory gene expression in L929sA versus TNFα alone.

**Conclusion:** GW enhances the anti-inflammatory effects of glucocorticoids in human FLS and L929sA, and mitigates metabolic side effects of dexamethasone in vivo, without compromising their efficacy. In addition, PPARα agonism permits to broaden its anti-inflammatory profile to interferon driven pathways. Given that both synthetic glucocorticoids and PPAR agonists are already widely used in (general) clinical practice, these findings offer a promising strategy to optimize glucocorticoid-based therapies.

**KEY MESSAGES:** *What is already known on this topic:*

- Synthetic glucocorticoids such as dexamethasone are widely used as immunosuppressive drugs, but cause many (metabolic) side effects.
- Peroxisome-proliferator-activated nuclear receptor (PPAR) agonists are clinically primarily used to treat symptoms that resemble glucocorticoid-induced side effects, but they also exert (modest) immunosuppressive effects.

*What this study adds:*

- This study explores the therapeutic potential of a combination treatment with dexamethasone and PPARα agonist GW7647 (GW) in cellular and murine models of rheumatoid arthritis
- We reveal that GW attenuates dexamethasone-induced adiposity and hypertriglyceridemia in vivo, hereby improving glucocorticoid-related side effects.
- The therapeutic efficacy of dexamethasone is maintained or even enhanced by GW in murine and cellular models of rheumatic arthritis, respectively.
- We offer novel mechanistic insights in the proposed combination treatment by revealing the selective suppression of interferon signaling pathways in human FLS.

*How this study might affect research, practice or policy:*

- Given that both synthetic glucocorticoids and PPAR agonists are already used in clinical practice, this study offers a promising, translatable strategy to optimize glucocorticoid-based therapies with an improved efficacy/safety ratio.

## INTRODUCTION

Up to 50% of rheumatoid arthritis (RA) patients take synthetic glucocorticoids. Despite their high efficacy, glucocorticoids can cause serious side effects such as diabetes, dyslipidemia and adiposity [1]. In RA, fibroblast-like synoviocytes (FLS), rather than immune cells, have been identified as the principal therapeutic targets of glucocorticoids [2].

Glucocorticoids act via the glucocorticoid receptor α (GRα, hereafter referred to as GR), a ligand-activated nuclear receptor and transcription factor that regulates gene expression assisted by coregulators. GR binds to glucocorticoid response elements (GREs) in promoter or enhancer regions of target genes as a homo(di)mer [3-5]. GR tethering to other transcription factors, such as AP1 and NF-κB, whose response elements are present in the promoters of pro-inflammatory genes, typically results in target gene suppression. Recently, a role for (atypical) heterodimers between GR and other nuclear receptors has been postulated, including peroxisome proliferator-activated receptors (PPARs) [6-9].

Like GR, PPARs suppress inflammation via NF-κB- and AP1-targeting mechanisms [10], while PPAR agonists are frequently used in the clinic to treat symptoms similar to glucocorticoid-related side effects, including prediabetes and dyslipidemia [11]. The three different PPAR isotypes are all activated by fatty acids and derivatives, or isotype-specific synthetic ligands, yet they regulate distinct aspects of metabolism [10, 12]. Interestingly, activation of both PPARα and PPARγ reduces arthritis severity in mouse and rat models, while PPARγ agonists also suppresses activation and inflammation in TNFα-induced FLS [13, 14].

Given the accumulating evidence that GR-PPAR receptor crosstalk can modulate both inflammatory and metabolic pathways [7, 15, 16], we explored whether combined GR and PPAR agonism could augment GR-mediated immunosuppression and/or attenuate its associated metabolic (side) effects in cellular and murine models of RA.

## MATERIALS AND METHODS

### Compounds and cytokines

Recombinant murine and human tumor necrosis alpha (TNFα) was produced and purified to 99% homogeneity at the VIB protein service facility. Recombinant mouse interferon-beta (IFNβ) was purchased at Biotechne (R&D Systems). Chicken collagen type II (CII) (MD Biosciences) was dissolved in 0.1M acetic acid at a concentration of 4 mg/ml by stirring overnight at 4 °C. Dexamethasone, GW7647 (GW) and WY-14643 (WY) were purchased from Sigma-Aldrich; rosiglitazone (ROSI) and GW501516 (GW50) from Cayman Chemicals.

### Collagen-induced arthritis mouse experiment

All animal procedures were approved by the institutional animal care and ethics committee (ECD23-89). 8-week old male DBA/1 mice were group-housed (5-8 mice/cage) in open cages, with a 12:12 light-dark cycle and *ad libitum* access to food and water (lights on at 7:00 AM, Zeitgeber time (ZT) = 0). To induce arthritis, mice were immunized with 200 µg chicken collagen type emulsified in Freund’s complete adjuvant (MD Biosciences), which was repeated after 3 and 5 weeks. At day 0, mice with a total arthritis severity score of >1 were selected and randomized based on total arthritis severity score, body weight and cage. Mice were allocated to the following treatment groups: Vehicle (3% DMSO in saline), GW (10 mg/kg/d), dexamethasone (0.25 mg/kg/d) and dexamethasone + GW (N=14/group); drug doses were determined based on previous findings [17-19]. From day 1, mice received daily drug injections i.p. between ZT2-ZT4 for 14 days. The clinical severity of arthritis per paw was graded daily according to standard evaluation procedures (see **supplementary materials**). At day (d)4, d7 and d14, body weight was measured. At d9, mice were fasted for 8h, after which an i.p glucose tolerance test was performed at ZT2. First, baseline samples were collected for measurements of plasma glucose (Freestyle, Freedom Lite), insulin (CrystalChem), triglycerides (Roche), and total cholesterol levels (Roche). Then, 2 mg/kg glucose was administered and blood samples were collected after 15, 30, 60 and 120min. At d14, mice were fasted for 8h. Mice received a final i.p. injection 1.5h before blood was collected retro-orbitally for VetScan HM5 (Bioscience, Zoetis) and IgG level analysis (in-house ELISA, see **supplementary materials**), after which mice were euthanized via cervical dislocation between ZT2-5. Mice were perfused with ice-cold PBS for 5min, after which tissues were isolated and stored until further processing.

### Cell Culture

#### Cell lines

All cells were maintained in a 5% CO2-humidified atmosphere at 37°C. Human FLS were grown in DMEM (Invitrogen) with 10% fetal calf serum (Invitrogen) and 1% Pen/Strep in a 5% CO2-humidified atmosphere at 37 °C. Murine fibrosarcoma L929sA cells were maintained in Dulbecco’s Modified Eagle Medium (DMEM, Invitrogen) plus 10% fetal calf serum. Murine fibrosarcoma L929sA cells with a stably integrated NFκB-driven reporter construct based on the E-selectin or IL-8 promoter [20], or a GRE-driven reporter construct [21] were cultured in DMEM plus 10% fetal calf serum and 500 µg/mL neomycin. The A549 and L929sA cell lines were verified to express GR, PPARα and PPARγ receptors endogenously [15].

### FLS experiments

The study was approved by the Ghent University Hospital Ethics Committee (BC-06496). Synovial tissues were obtained during joint replacement surgery from four female osteoarthritis patients with BMI >25, who were not using any GR, PPARα or PPARy agonists. Informed consent was obtained from all participating patients. FLS were obtained by enzymatic digestion as previously described [22]. Experiments were performed using FLS with a passage number ranging from 4 to maximally 8. FLS were seeded in three technical replicates in 12-well plates (100,000 cells/well) and, 24h later, were serum deprived for 16h prior to compound stimulations for 6h for transcriptional analyses, and 24h for protein and secretome analyses. Subsequent procedures are described in the **supplementary materials**.

### Cell reporter assays

L929sA reporter cells (driven by GREs or E-selectin/IL8 promoters) were seeded in black clear bottom 96-well plates (5000cells/well). After 24h, cells were serum-deprived for 16h prior to compound stimulations. Cells were lysed with 1x Cell Culture Lysis Reagent (Promega) and cell lysates were assayed for luciferase activity (EnVision 2102 Multilabel reader, Perkin Elmer) by addition of luciferin substrate (manufactured as described previously [23]), and normalized for beta-galactosidase activity (Galacto-Star™, ThermoFisher Scientific).

### Statistical analysis

Statistical analyses were performed with GraphPad Software (version) and R (version 4.3.2.). All data are expressed as mean ± SEM. All p-values are two-tailed and p<0.05 was considered as statistically significant. Data with one, two (i.e. dexamethasone and GW) or three factors (i.e. dexamethasone, GW, and time/IFNAR-ab) were analyzed with a one-way, two-way or three-way ANOVA respectively, and a Tukey post-hoc test. Figures show post-hoc comparisons made between 1) TNF-stimulated vehicle and all other groups and 2) dexamethasone with and without GW. When 2-way ANOVA analysis was performed, graphs additionally show the statistical significance of all main effects below the graph (D = Dexamethasone, G = GW, I = Dexamethasone-GW interaction). To calculate statistical significance for concentration-response curves in luciferase reporter assays, gamma generalized linear models were performed with averaged pseudoreplicates, with respective compounds and experiment:plate effects as variables. Details on the proteomics data analyses can be found in the **supplementary materials**.

### A Patient and Public Involvement

The conceptual basis of this study was disseminated at ‘Reumacafé’ organized by the Flemish patient society for rheumatic diseases (ReumaNET). Patients and/or the public were not further involved in the design, conduct or reporting of this research.

## RESULTS

### Dexamethasone and GW7647 cooperatively suppress TNFα signaling in murine L929sA fibroblasts

We first determined the optimal concentrations of the GR agonist dexamethasone and the PPARα agonist GW7647 (GW) to reveal potential cooperative immunosuppressive effects. L929sA served as a first ‘workhorse’ model, because they express both GR and PPARα [15] and, being fibroblasts, share many characteristics with fibroblast-like synoviocytes (FLS) [24]. L929sA cells containing a stably integrated, TNFα-responsive, E-selectin- or IL-8-promoter based luciferase reporter construct were pre-treated with increasing concentrations of dexamethasone with and without GW for 1h, before stimulation with TNFα for an additional 5h. TNFα strongly upregulated E-selectin and IL-8 promoter activity (200-fold and 22-fold, respectively, **Sup. Fig. 1-B**). Dexamethasone alone concentration-dependently reduced E-selectin and IL-8 activity (**Fig. 1A-B**). Interestingly, this reduction was markedly enhanced in the presence of GW (**Fig. 1A-B**). In a similar set-up, E-selectin/IL-8 reporter cells were treated with increasing concentrations of GW, with and without dexamethasone. GW alone suppressed reporter E-selectin and IL-8 activities at relatively high concentrations (≥0.5μM, **Fig. 1C-D**), and with the greatest efficacy in the E-selectin reporter cell line. Again, the combination of GW and dexamethasone suppressed inflammatory activities more strongly compared to either drug alone (**Fig. 1A-D**). For subsequent experiments, we selected drug concentrations that suppressed TNFα signaling by 50-65%, which is 0.01µM for dexamethasone and max 2.5μM for GW.

**Figure 1.**
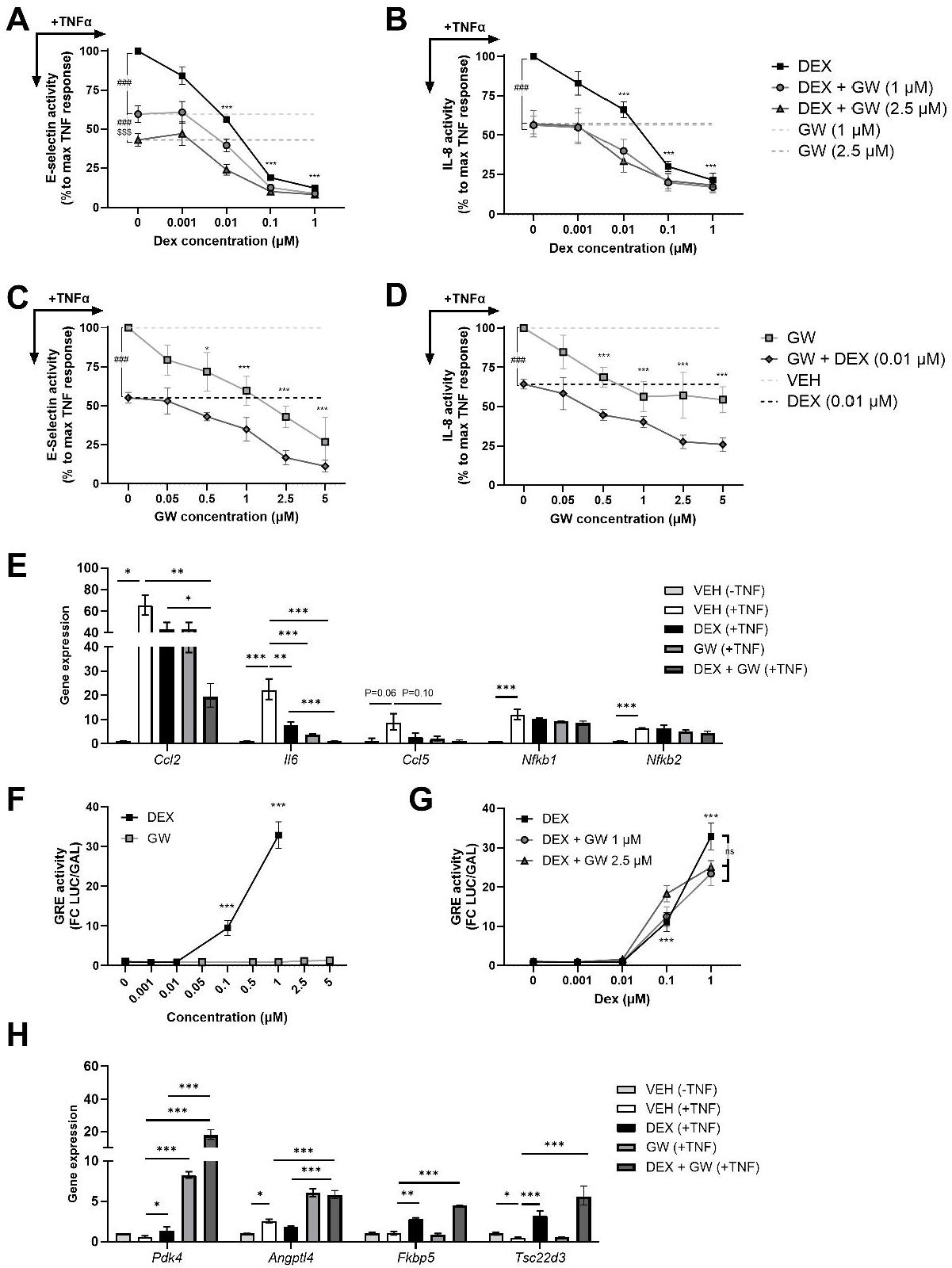
Dexamethasone and GW7647 cooperatively suppress TNFα signaling while increasing expression of (shared) GR/PPAR target genes in murine connective tissue L929sA cells. A-D) L929sA cells containing a stably integrated, TNFα-responsive E-selectin- or IL-8-based promoter luciferase reporter construct were pre-treated with (combinations of) dexamethasone (DEX), GW7647 (GW) or vehicle (VEH) for 1h, before stimulation with murine TNFα for an additional 5h. Cell lysates were assayed for luciferase activities and normalized for β-galactosidase activities. E) Expression of pro-inflammatory genes after (combinations of) 0.01μM DEX and 2.5μM GW, measured by qPCR and shown as fold change relative to the TNF(-) group. F-G) L929sA cells containing a stably integrated glucocorticoid-responsive GRE-reporter construct were treated with (combinations of) DEX, GW or vehicle (VEH) for 6h, after which luciferase activity was evaluated. H) L929sA cells were pre-treated with (combinations of) dexamethasone (0.01μM, DEX), GW7647 (2.5μM, GW) or vehicle (VEH) for 1h before stimulation with TNFα (+) or medium (-) for an additional 5h, after which expression of (shared) GR and PPAR target genes was measured by qPCR and shown as fold change relative to the TNF(-) group. Statistical significance was calculated by gamma GLM in A-D) and F-G); by one-way ANOVA with Tukey multiple comparison tests in E) and H) on N=3 experiments. Results are shown as mean ± SEM. *P<0.05; **P<0.01; ***P<0.001; ###P<0.001 vs DEX; $$$P<0.001 vs GW 1 µM.

As a complementary approach, we investigated immunosuppression in an endogenous gene regulation context, by analyzing dexamethasone and GW effects on pro-inflammatory gene transcripts in TNFα-stimulated L929sA. TNFα robustly increased the expression of *Ccl2, Il6, Ccl5, Nfkb1* and *Nfkb2* (**Fig. 1E**). Treatment with dexamethasone (0.01µM) or GW (2.5µM) alone reduced the expression of *Ccl2, Il6, Ccl5*, but had no effect on *Nfkb1* and *Nfkb2*. Notably, combined dexamethasone and GW treatment significantly enhanced the suppression of TNFα-induced *Ccl2* and *Il6* expression compared to dexamethasone alone, yet only a trend was observed for *Ccl5* (P=0.10, **Fig. 1E**). Altogether, these findings suggest that a subset of TNFα-regulated pro-inflammatory genes is more effectively suppressed by combined dexamethasone and GW treatment than by either compound alone in L929sA fibroblasts.

### Dexamethasone and GW7647 cooperatively regulate shared target genes

We subsequently studied crosstalk mechanisms between GR and PPARα at the level of transcriptional activation (reporter assays) and (shared) GR and PPARα target gene regulation (qPCR) upon dexamethasone and/or GW treatment of L929sA cells. As expected, dexamethasone, but not GW, concentration-dependently increased GRE-dependent reporter activity in L929sA cells (**Fig. 1F**). Only a trend was noted for GW-mediated reduction of GRE activity when combined with high concentrations of dexamethasone (Main GW effect P=0.08, (**Fig. 1G**), indicative for a mild crosstalk effect. At the endogenous level, dexamethasone (0.01µM) alone upregulated the expression of most studied genes (*Pdk4, Fkbp5* and *Tsc22d3*), while GW alone (2.5µM) only increased *Pdk4* and *Angptl4* expression in L929sAs (**Fig. 1H**). Interestingly, GW markedly enhanced the dexamethasone-induced upregulation of both *Pdk4* and *Angptl4* – highlighting the presence of GR-PPAR crosstalk through the cooperative regulation of shared target genes.

### GW7647 reduces dexamethasone-induced hypertriglyceridemia and adiposity in collagen-induced arthritis mice

Encouraged by the enhanced anti-inflammatory effects in the L929sA cell model, we investigated the metabolic and immunosuppressive effects of dexamethasone and GW in a collagen-induced arthritis (CIA) mouse model, using a relatively low dexamethasone dose (0.25 mg/kg/d [18]) to allow detection of potential additive immunosuppressive GW effects. Dexamethasone reduced body weight when corrected for body weight at start, but not when uncorrected (main effect ANOVA significant, **Fig. 2A-B**). GW addition did not alter these observations. Dexamethasone increased gonadal white adipose tissue, which was partially prevented by GW (main effect ANOVA significant, **Fig. 2C**). GW increased liver weight, which was reduced by adding dexamethasone (**Fig. 2D**). Regarding glucose metabolism, dexamethasone induced hyperinsulinemia, which was not influenced by GW (**Fig. 2E**). In addition, none of the treatments influenced basal glucose levels nor the clearance of glucose after an intraperitoneal glucose injection (**Fig. 2F-H**). Regarding lipid metabolism, GW alone reduced plasma triglyceride levels whereas dexamethasone alone induced hypertriglyceridemia (**Fig. 2I**). Notably, the dexamethasone-induced hypertriglyceridemia was fully prevented when GW was combined with dexamethasone (**Fig. 2I**). Finally, dexamethasone slightly increased total cholesterol levels, which were not affected by GW (main effect ANOVA significant, **Fig. 2J**). In conclusion, GW reduced adiposity and hypertriglyceridemia induced by dexamethasone, but not the hyperinsulinemia or hypercholesterolemia in the CIA model - consistent with well-described effects of PPARα agonists.

**Figure 2.**
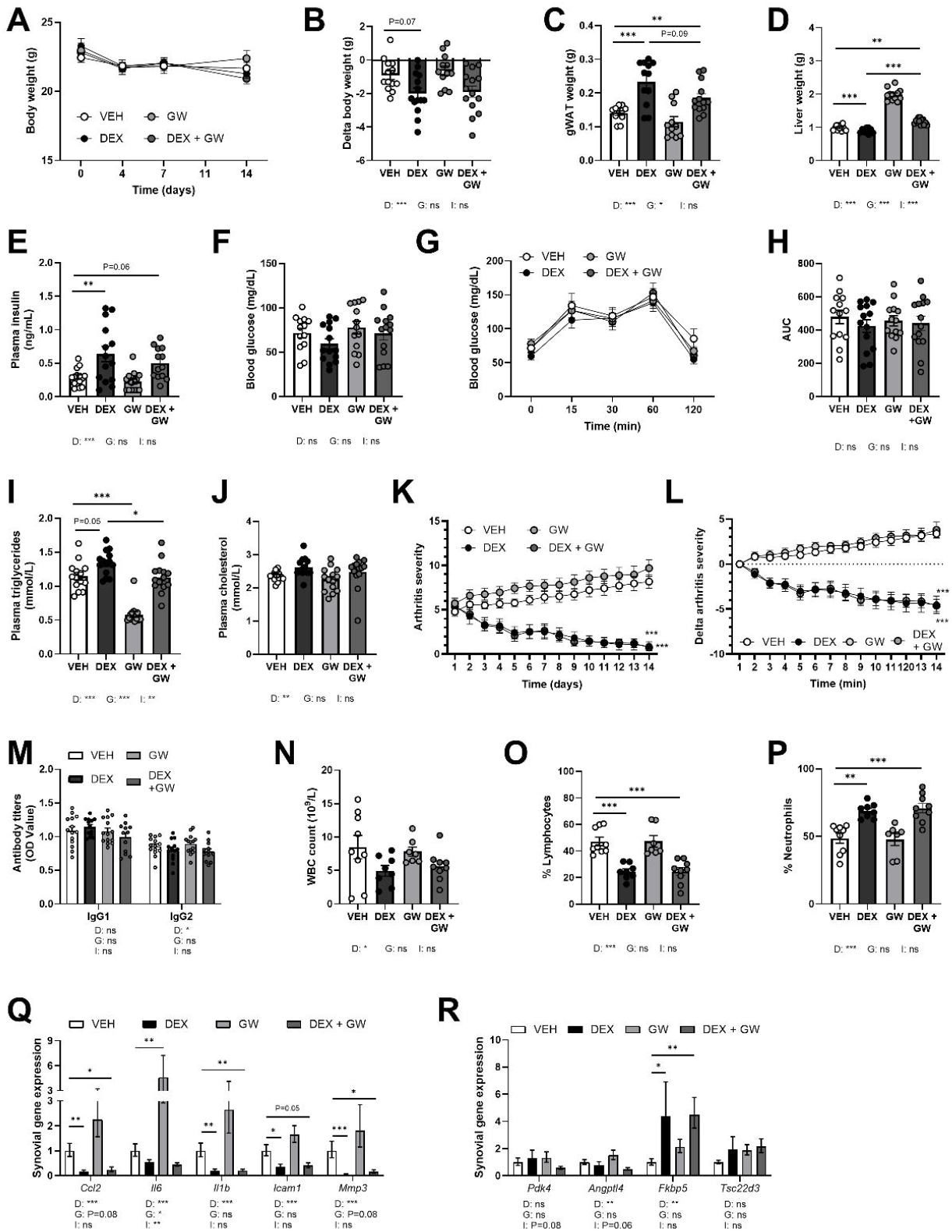
GW7647 (partially) reduces hypertriglyceridemia and adiposity induced by dexamethasone in collagen-induced arthritis mice. CIA mice (N=14/group) were treated with (combinations of) vehicle (VEH), GW7647 (GW, 10 mg/kg/d) and dexamethasone (DEX, 0.25 mg/kg/d) for 15 days. A-B) Body weight was measured across the experiment, raw values expressed in A), and in B) corrected for start body weight. C-D) Wet tissue weights of gonadal white adipose tissue (gWAT) and liver. E-J) On day 10, mice were fasted for 8h, blood was collected for insulin, glucose, triglyceride and total cholesterol measurements, followed by an intraperitoneal glucose tolerance test. H) The area under the curve (AUC) was calculated from the individual glucose curves. K-L) Arthritis severity was scored across the experiment, raw values expressed in K), and in L) corrected for start severity score. M) Antibody (IgG1 and IgG2) titers on day 15. N-P) Total white blood cell (WBC) counts and other immune populations on day 15. Q) Expression of inflammatory genes in knee synovia and R) expression of (shared) GR and PPAR target genes in knee synovia was measured by qPCR and shown as fold change relative to the TNF(-) group. Statistical significance was calculated by three-way ANOVA for A); two-way ANOVA for B-R). Results are shown as mean ± SEM. Main effects from 2-way ANOVA analyses are depicted under graphs (D = DEX, G = GW, I = DEX-GW interaction). Post-hoc comparisons are shown in the figure as *P<0.05; **P<0.01; ***P<0.001.

### GW7647 did not enhance dexamethasone’s ability to reduce arthritis severity in collagen-induced arthritis mice

With regard to anti-inflammatory effects, dexamethasone gradually reduced the mean arthritis severity score over several days, which was not influenced by the addition of GW (**Fig. 2K-L**). GW alone slightly increased the mean arthritis severity score, although this effect was not significant and fully abolished when corrected for baseline severity (**Fig. 2K-L**). Dexamethasone, but not GW, slightly decreased plasma anti-collagen IgG2 levels (main effect ANOVA significant, **Fig. 2M**), with no effect on IgG1. Dexamethasone, regardless of the presence of GW, decreased total white blood cell count and percentage lymphocytes, but increased percentage neutrophils (**Fig. 2N-P**). Dexamethasone’s immunosuppressive actions were confirmed with complementary gene expression analyses, as we found reduced expression of TNFα-regulated genes implicated in RA pathology in knee synovial tissue (**Fig. 2 Q**) [25]. Unexpectedly, and in contrast to our *in vitro* data in L929sA cells (**Fig. 1E**), GW alone upregulated the expression of *Il6*, and non-significantly of *Ccl2* and *Mmp3* (P=0.08 for both); effects that were fully prevented by dexamethasone (**Fig. 2Q**). Dexamethasone, in the presence or absence of GW, significantly induced *Fkbp5* expression. However, the (cooperative) effects of dexamethasone and GW on *Pdk4* and, *Angptl4* expression, as previously observed in L929sAs, were not recapitulated in mouse synovia (**Fig. 1H, Fig. 4R**).

### PPARα protein levels are markedly reduced upon combined dexamethasone and GW7647 treatment in serum-starved, human fibroblast-like synoviocytes

Intrigued by the context-specificity of the cooperative effects of dexamethasone and GW, we aimed to dissect the molecular basis of this crosstalk in a model that more closely mimics RA-pathology: human FLS, isolated from synovial tissues collected during joint replacement surgery from four female osteoarthritis patients with BMI >25, who were not using any GR or PPAR agonists. FLS from all four patients expressed GR and PPARα, and downregulated GR protein levels only in response to dexamethasone (**Fig. 3A**). Interestingly, we found that PPARα levels were markedly reduced only after combined dexamethasone and GW treatment. In a pilot gene expression analysis experiment, we observed that serum-starvation enhanced the effects of combined dexamethasone and GW treatment on *ANGPTL4* and *FKBP5* expression, but not on *PDK4* expression (**Sup. Fig. 2A-C**). Dexamethasone and GW alone upregulated *PDK4* and *ANGPTL4* expression, which was markedly enhanced by combined dexamethasone and GW treatment (**Sup. Fig. 2A-B**). As serum-starvation also enlarged the TNFα-induced expression of pro-inflammatory genes *CCL2, IL6* and *CCL5* (**Sup. Fig. 3A-C**), we decided to proceed with serum-starvation for subsequent experiments.

**Figure 3.**
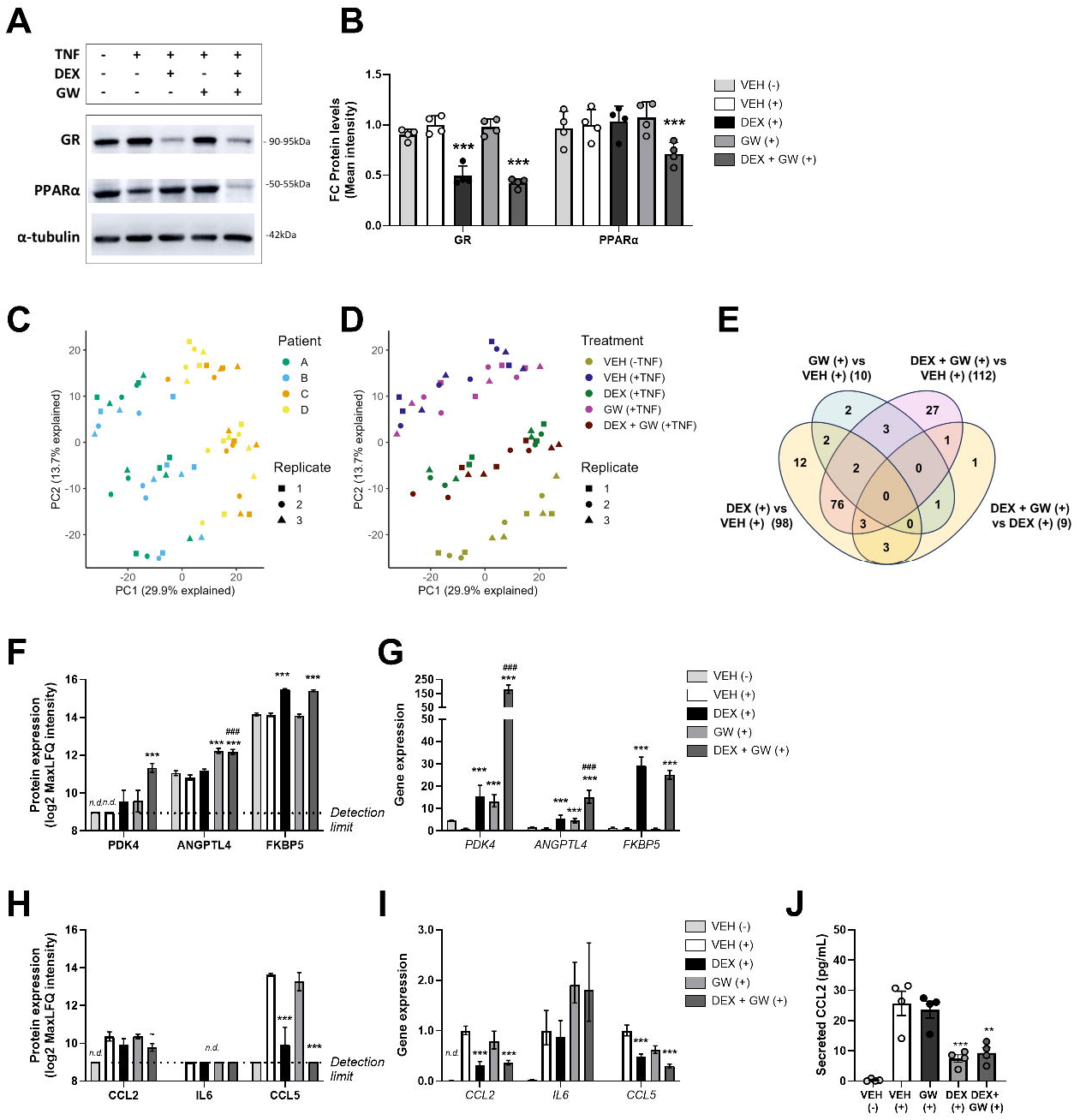
GR-PPAR crosstalk is observed in human fibroblast-like synoviocytes. Human fibroblast-like synoviocytes (FLS) were serum-starved for 16h before the experiment. Cells were pre-treated with (combinations of) dexamethasone (0.01μM, DEX), GW7647 (2.5μM, GW) or vehicle (VEH) for 1h, before stimulation with human TNFα (+) or medium (-) for an additional 23h for A-B) Western Blot analysis, C-F, H) shotgun proteomics; and supernatant analysis (J). TNFα stimulation was only 5h for transcriptional analyses in G) and I) using qPCR. A) Representative Western Blot of FLS from one patient. B) Quantified FLS protein levels from Western Blots (N=4 patients). C-D) Principal component analyses of FLS proteome based on quantified protein expression values (N=4 patients). E) Venn diagram showing the number of significantly differentially expressed proteins. F-G) Expression of GR and PPAR target proteins and genes in FLS, respectively. H-I) Expression of pro-inflammatory proteins and genes, respectively. Gene expression was measured by qPCR and shown as fold change relative to the TNF(+) group. J) Secreted CCL2 levels in supernatant. Statistical significance was calculated on N=4 patients by one-way ANOVA and Tukey’s post-hoc test in A-B), G) and I-J); and by the package limma in E-F) and H), using a false discovery rate (FDR) of < 0.05, a fold change of ≥ 2- or ≤ 0.5-fold (|log2FC| ≥ 1) and an adjusted p-value < 0.05. Results are shown as mean ± SEM. n.d. = not detected. *P<0.05; **P<0.01; ***P<0.001; ###P<0.001 vs DEX.

### TNFα and dexamethasone induce the largest divergence in fibroblast-like synoviocytes’ proteome profiles

To evaluate the broader impact of dexamethasone and GW (combination) treatments, we performed proteomics in TNFα-stimulated, serum-starved FLS upon 24h-compound stimulation. We detected 7703 protein groups in all samples, of which 7686 were reliably quantified (**supplementary materials**). PCA analysis revealed that most variation was explained by patient origin (PC1, **Fig. 3C**), while TNFα and dexamethasone treatments caused the largest separation of clusters (PC2, **Fig. 3D**). In line with the latter finding, samples treated with GW clustered together with TNFα-stimulated samples, while samples treated with dexamethasone, with and without GW, clustered together (**Fig. 3D**) – indicating that the FLS proteome was much stronger affected by dexamethasone and TNFα than by GW. We next conducted a differential expression analysis comparing dexamethasone, GW and combined dexamethasone and GW treatments with the TNFα-stimulated vehicle group, and dexamethasone alone versus dexamethasone plus GW, using an adjusted p-value < 0.05 and a log-fold change threshold (|log2FC| ≥ 1). Dexamethasone alone and dexamethasone plus GW differentially regulated 17 and 31 proteins, respectively (**Fig. 3E**), although most differentially regulated proteins were shared between the two groups (81 overlapping proteins). GW alone significantly regulated only 10 proteins, of which 2 uniquely (**Fig. 3E**). Finally, when comparing the dexamethasone versus dexamethasone plus GW treated groups, only 9 proteins were statistically different (**Fig. 3E**).

### GR-PPAR crosstalk is observed in human fibroblast-like synoviocytes

As GW regulated relatively few proteins in FLS, we investigated whether the regulation of (shared) GR and PPAR targets and inflammatory targets, previously observed in L929sA, was replicated in FLS. Dexamethasone and GW each increased PDK4 protein expression, and this increase was substantially larger after combined treatment (**Fig. 3F**). Similar findings were observed at the transcriptional level, which completely aligns with the previously observed crosstalk (**Fig. 3G, Fig. 1H**). In the case of ANGPTL4, crosstalk between dexamethasone and GW was more apparent at the mRNA level than at the protein level. Finally, FKBP5 protein and gene expression aligned and were increased by dexamethasone, with and without GW (**Fig. 3F-G**). Regarding pro-inflammatory targets, modest crosstalk effects of dexamethasone and GW were observed. TNFα increased protein and mRNA levels of CCL2 and CCL5, but only at the mRNA level for IL6 (**Fig. 3H-I**). Dexamethasone combined with GW reduced intracellular CCL2 protein (P=0.003, log2FC=- 0.63) and mRNA levels (**Fig. 3H-I**), and also secreted CCL2 protein levels, although these effects were not different from dexamethasone treatment alone (**Fig. 3I-J**). Combined dexamethasone and GW treatment, however, did suppress TNFα-induced CCL5 protein and gene expression to a somewhat higher extent than dexamethasone alone (**Fig. 3H-I**). In conclusion, GR-PPAR crosstalk effects were observed at the mRNA and protein level for PDK4 and to a minor extent for CCL5 and ANGPTL4 in FLS, suggesting target-specific regulation.

### GW7647 enhances suppression of interferon-stimulated proteins after dexamethasone in human fibroblast-like synoviocytes

Besides the hypothesis-driven interrogation of specific protein targets (**Fig. 3F-J**), we performed protein set enrichment analysis, comparing proteins differentially expressed upon dexamethasone plus GW versus dexamethasone alone. Given the limited number of significantly regulated proteins (**Fig. 3E**) and the integration of log_2_ fold change information within Ingenuity Pathway Analysis (IPA), we performed IPA analysis on all proteins with FDR-adjusted p<0.05, but without applying an additional |log_2_FC|≥1 threshold, leading to N=82 proteins as input for IPA. Surprisingly, among the top-ranked canonical pathways identified, the first, third, and fourth were related to interferon signaling (**Fig. 4A**). Network analysis on interactions between differentially expressed proteins and IPA-predicted effects also revealed many well-described interferon-stimulated proteins (ISPs), which had TNFα, NF-κB and IFNα/β as predicted suppressed regulators (**Fig. 4B**). Subsequently, all ISPs identified in the dataset were selected for heatmap visualization and hierarchical clustering (**Sup. Fig. 4**). This revealed a subset of 38 ISPs which were more strongly downregulated after combined dexamethasone and GW treatment compared to dexamethasone alone, including MX1, HERC6, IFI44, IFI44L, ISG15 and USP18 - which showed the most striking differential response among the subset – and STAT1 and STAT2, interesting in the light of the recent introduction of JAK/STAT inhibitors as RA treatment (**Fig. 4C, Sup. Fig. 4**). On the individual protein level, MX1 and HERC6 proteins were downregulated by dexamethasone, and potentiated further by GW (P<0.05, |log_2_FC| ≥ 1, **Fig. 4C**). IFI44, IFI44L and ISG15 were also all downregulated by dexamethasone, and more so after combined dexamethasone and GW treatment, although to a lower extent as MX1 and HERC6 (P<0.05, but |log_2_FC| < 1; **Fig. 4C**). The dexamethasone and GW crosstalk was less pronounced at the transcriptional level for these targets (**Fig. 4D**). Taken together, GW selectively potentiated the dexamethasone-mediated suppression of interferon-related signaling in human FLS.

**Figure 4.**
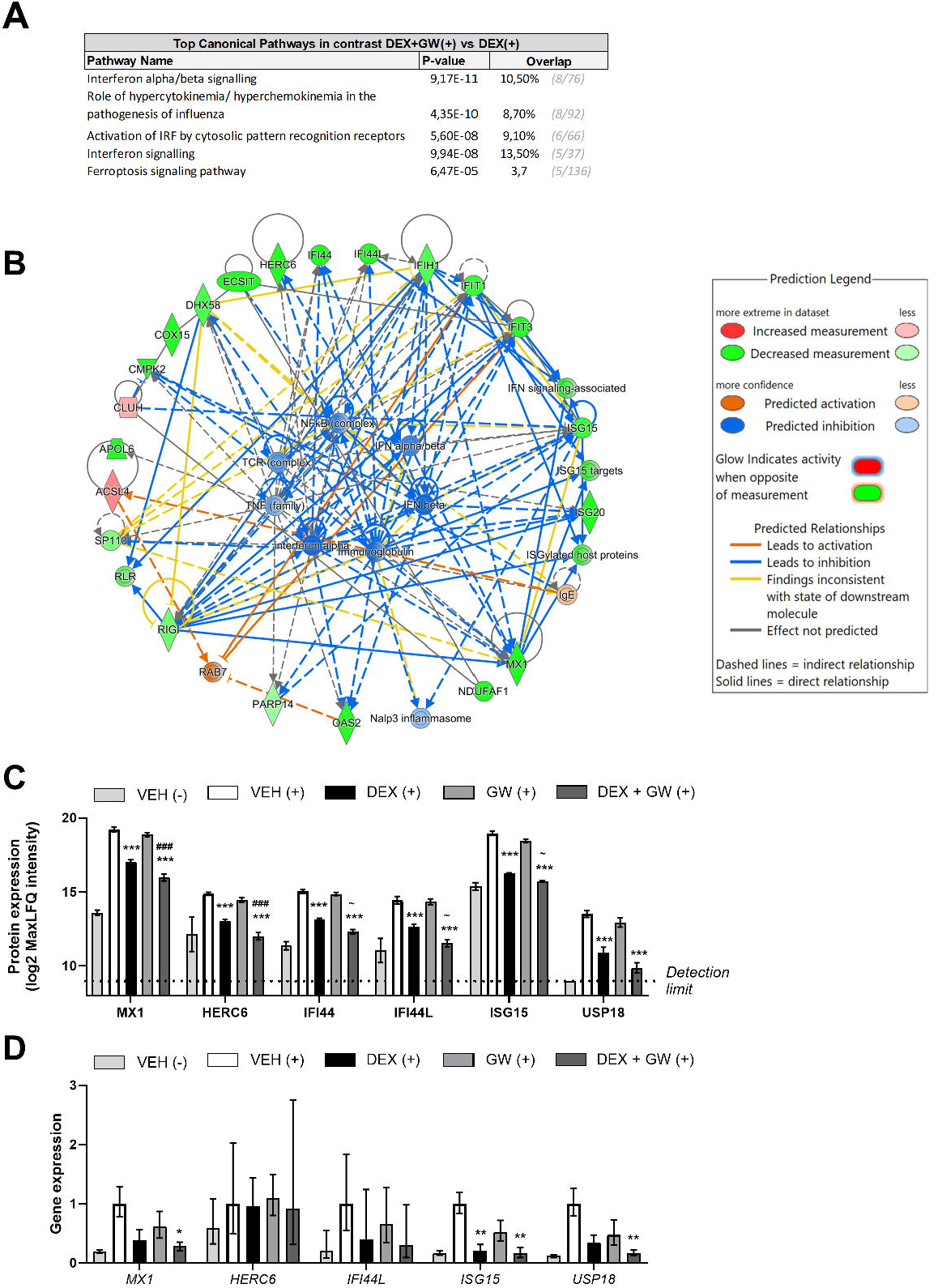
Interferon-stimulated proteins are more downregulated by combined dexamethasone and GW7647 treatment. Human fibroblast-like synoviocytes (FLS) were serum-starved for 16h before the experiment. Cells were pre-treated with (combinations of) dexamethasone (0.01μM, DEX), GW7647 (2.5μM, GW) or vehicle (VEH) for 1h, before stimulation with human TNFα (+) or medium (-) for an additional 23h, followed by shotgun proteomics. A) Top canonical pathways found in the DEX+GW vs DEX contrast by Ingenuity Pathway Analysis (IPA). B) Network analysis revealing interactions between differentially expressed proteins and IPA-predicted effects. C-D) Expression of interferon-regulated proteins and genes, respectively; gene expression was measured by qPCR and shown as fold change relative to the TNF(+) group. Statistical significance was calculated by differential expression analysis on N=4 patients using package limma, with a false discovery rate and adjusted p-value < 0.05 for A-B), and additionally a fold change of ≥ 2- or ≤ 0.5-fold (|log2FC| ≥ 1) for C). For D), statistical significance was calculated with one-way ANOVA. Results are shown as mean ± SEM. *P<0.05; **P<0.01; ***P<0.001 vs VEH. ∼P<0.05 vs VEH, but did not reach the |log2FC| ≥ 2- or ≤ 0.5 threshold; ###P<0.001 vs DEX

### Cooperative immunosuppressive effects of dexamethasone and GW are not mediated through inhibition of autocrine IFNβ signaling

Because we found a unexpected dependency on interferon-related signaling in our shotgun proteomics - and because TNFα can stimulate autocrine IFNβ production in both L929sA and FLS [26, 27] - we pre-treated L929sA with an antibody blocking the IFNα/β receptor (IFNAR), followed by stimulation with TNFα and (combined) dexamethasone and GW treatment. We did not find evidence that the cooperative anti-inflammatory effects of dexamethasone and GW were mediated via suppression of autocrine IFNβ production, as the IFNAR-antibody did not influence the suppression of TNFα-regulated genes *Ccl2* and *Ccl5* by dexamethasone and GW (**Fig. 5A-B**). We next asked whether conversely, the cooperative immunosuppressive effects of dexamethasone and GW may become more evident after co-stimulation with TNFα and IFNβ. The expression of *Ccl5*, but not *Ccl2*, was 10-fold increased by combined stimulation of TNFα and IFNβ compared to TNFα alone, which could largely be prevented by IFNAR blockade (**Fig. 5C-D** vs. **5A-B**). Interestingly, combined dexamethasone and GW treatment suppressed *Ccl5* much stronger after combined TNFα and IFNβ stimulation, than after TNFα alone, indicating that a vast suppression depends on the presence of both cytokines (**Fig. 5C**). In conclusion, the suppressive effects of combined dexamethasone and GW on inflammatory chemokines are both gene- and context-specific, with *Ccl5* suppression being more outspoken after co-stimulation of TNFα and IFNβ than after TNFα alone.

**Figure 5.**
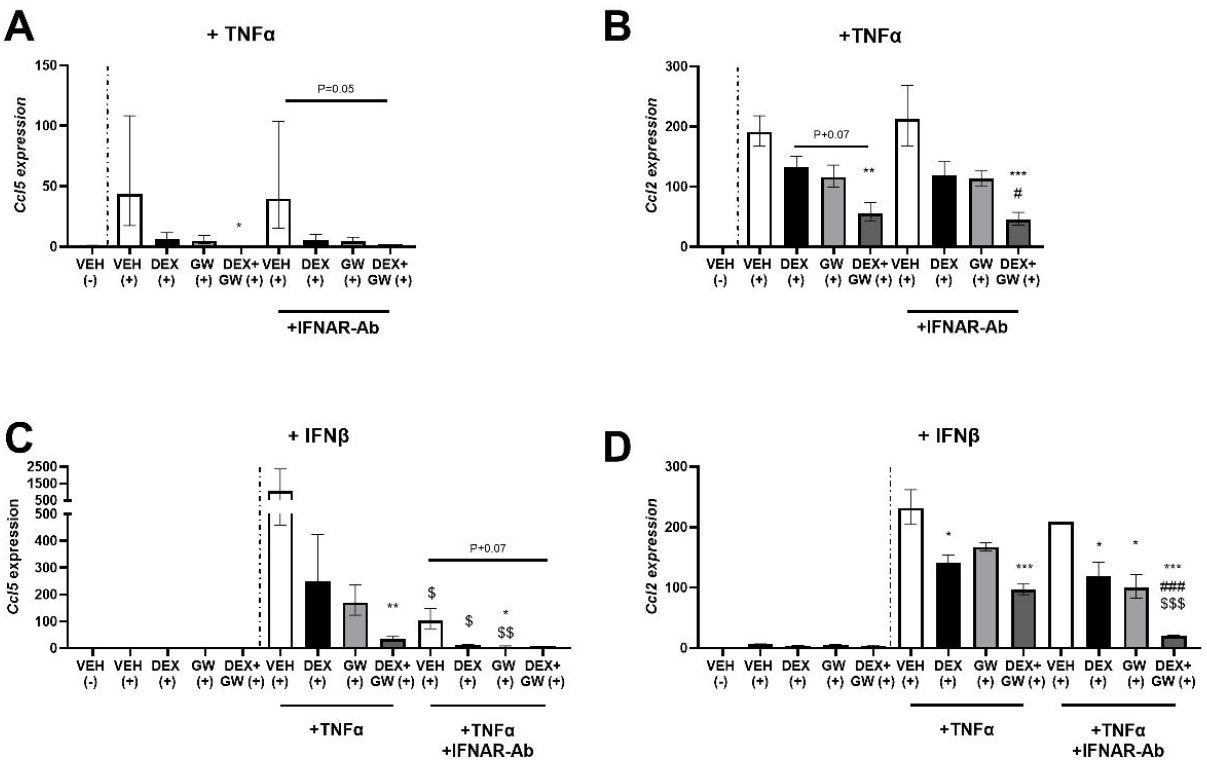
TNFα and IFNβ cooperatively increase *Ccl5* expression. L929sA cells were pre-treated with an IFNα/β receptor antibody (IFNAR-Ab) for 16h. Cells were subsequently treated with (combinations of) dexamethasone (0.01μM, DEX), GW7647 (2.5μM, GW) or vehicle (VEH) for 1h before stimulation with murine TNFα or IFNβ for an additional 5h, after which expression of pro-inflammatory genes was measured by qPCR and shown as fold change relative to the TNF(-) group. Statistical significance was calculated by three-way ANOVA on N=2-3 experiments. Results are shown as mean ± SEM. *P<0.05; **P<0.01; ***P<0.001 vs VEH. #P<0.05; ##P<0.01; ###P<0.001 vs DEX; $$$P<0.001 vs non-IFNAR-ab.

## DISCUSSION

This study explored whether GW could enhance the immunosuppression and/or mitigate any of the metabolic side effects of dexamethasone. We found that GW reduced the dexamethasone-induced hypertriglyceridemia and adiposity in CIA mice. GW potentiated the immunosuppressive effects of dexamethasone in TNFα-stimulated L929sA and human FLS, but not in CIA mice. Notably, combined dexamethasone and GW treatment target-selectively suppressed interferon signaling in TNFα-stimulated human FLS. In line with this, combined dexamethasone and GW treatment abrogated synergistic *Ccl5* expression following TNFα and IFNβ co-stimulation – suggesting that PPARα agonism can broaden the anti-inflammatory profile to interferon driven pathways.

In our CIA model, GW, with and without dexamethasone, did not reduce arthritis severity, contrasting with previous studies where the PPARα agonist fenofibrate and the PPARγ agonist rosiglitazone attenuated arthritis severity [13, 14]. Next to compound-specific pharmacokinetics- or dynamics, species- or model-specific differences may underlie this divergent outcome, as human RA is primarily driven by dysregulated adaptive immunity— including IFN signaling—whereas rodent CIA involves both adaptive and systemic innate immune responses [28]. Notably, strong IFNAR activation was previously observed in human FLS from RA patients, while this was absent in CIA rat paws [29]. Given that the cooperative immunosuppressive effects of dexamethasone and GW were most pronounced after TNFα/IFNβ co-stimulation, the lack of IFNAR pathway activation in CIA may partly explain the absence of synergistic immunosuppressive effects in our CIA model.

The cooperative suppression of interferon signaling by dexamethasone and GW in human FLS has substantial clinical relevance. Notably, the suppressed ISPs include key components of the ‘type I interferon signature’, such as MX1, IFL44L, ISG15 and OAS1 (**Fig. 4, Sup. Fig. 4**) [30]. An elevated baseline expression of the IFN signature in peripheral blood mononuclear cells predicts poorer 6-month outcomes and reduced responsiveness to disease-modifying antirheumatic drugs (DMARDs) in (early) RA patients [31-34]. Beyond driving disease progression, this interferon signature is also associated with an increased risk of developing RA in preclinical individuals [35]. In FLS specifically, TNFα stimulates autocrine IFNβ production through IRF1, which accounts for approximately 50% of the TNFα-driven transcriptional response [27, 36]. While we could not find evidence for autocrine IFNβ production in L929sA, it likely contributed to the response observed in FLS. Finally, reduced STAT1 and STAT2 levels after combined dexamethasone and GW treatment are noteworthy (**Sup. Fig. 4** ), given the introduction of JAK/STAT inhibitors [37], further supporting the therapeutic potential of targeting IFN signaling in RA.

In contrast to previous findings [15], GW did not significantly reduce dexamethasone-induced GRE-reporter activity or expression of the GR target gene *Gilz*. In this study, PPARα was not overexpressed, which may have diminished the GW effect. Additionally, the dexamethasone concentration was 10-to 100-fold lower in this study, while the attenuation of *Gilz* expression is concentration-dependent [15]. Finally, the different cellular context, i.e. L929sA versus HepG2 cells, may also contribute to this difference, as a different cellular repertoire of coregulators can drastically change the transcriptional response [15, 38].

GW reduced dexamethasone-induced hypertriglyceridemia and adiposity in CIA mice, but we were unable to corroborate in this CIA model our previous finding that PPAR agonism prevents dexamethasone-induced glucose intolerance. Indeed, dexamethasone did not induce glucose intolerance in the current study, possibly due to the lower dexamethasone dose and absence of high-fat diet feeding, as our main objective was to investigate potential cooperative effects on immunosuppression, and not metabolism [15]. Consistent with prior studies, we additionally observed GW increased liver weight, an effect shown to be fully PPARα-dependent [39]. Interestingly, the GW-induced increase in liver weight was prevented by dexamethasone, suggesting an additional layer of GR-PPAR crosstalk with potential therapeutic relevance.

Overall, GW attenuates some of the metabolic side effects of glucocorticoids, while maintaining (CIA) or even enhancing its therapeutic efficacy, especially in the context of interferon-driven pathways (human FLS, L929sA). Our findings are not only relevant for RA, but also for other arthritic conditions where (systemic) glucocorticoids remain essential and IFN-driven pathology is prominent. As both GR and PPAR agonists are already in widespread clinical use, their combination offers a strong and affordable translational potential to optimize therapeutic efficacy while minimizing side effects.

## Author contributions

Conceptualization and design: LK, DC, DE, KDB; Animal experiments and data analysis: LK, EG, JC; Cell line experiments and analysis: LK, AV, JT, MK; FLS experiments: LK, EG, JC, AV, GP; Proteomics analysis: LK, DF, SD, SdV; Statistical analysis: LK, LM; Manuscript original draft: LK; Manuscript reviewing and editing: EG, DC, DF, SD, LM, DE, KDB; All authors reviewed and approved the manuscript.

## Acknowledgements

We thank Delphi Van Haver (VIB proteomics core) and Trea Streefland (LUMC) for their valuable technical support. Additionally, we are grateful to Edmee Vandenboorn (TU Eindhoven) for performing valuable pilot experiments, and to Eleni Staessens (VIB-UGent) for her assistance with the FLS experiments.

