## supplementary materials for "Optimizing efficacy to safety ratio of glucocorticoids in rheumatoid arthritis models by leveraging PPARα agonism"

[*]*Shared authorships*

[1]*VIB Center for Medical Biotechnology, Department of Biomolecular Medicine, Ghent University, Ghent, Belgium;*
[2]*Department of Anatomy and Embryology, Leiden University Medical Center, Leiden, Netherlands*
[3]*VIB Center for Inflammation Research, Ghent University and Department of Rheumatology, Ghent University Hospital, Ghent, Belgium.*
[4]*Cancer Research Institute Ghent (CRIG), Corneel Heymanslaan 10, 9000 Gent, Belgium.*
[5]*VIB Proteomics Core, Ghent, Belgium*
[6]*Department of Orthopaedics and Traumatology, Ghent University Hospital, Ghent, Belgium.*
[7]*VIB Center for Medical Biotechnology, Department of Biochemistry and Microbiology, Ghent University, Ghent, Belgium*

### SUPPLEMENTARY FIGURES

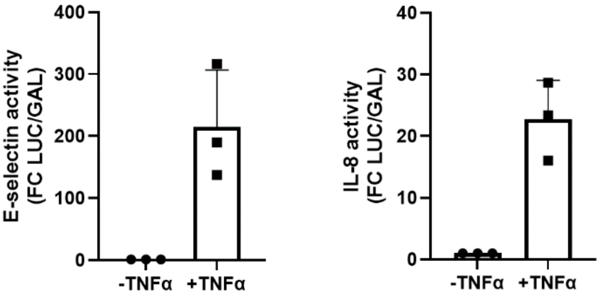


**Supplementary figure 1 - TNFα signaling strongly increases E-selectin and IL-8-reporter activity in murine connective tissue L929sA cells.** A-B) L929sA cells containing a stably integrated, TNFα-responsive E-selectin- or IL-8-based promoter luciferase reporter construct were stimulated with murine TNFα or medium for 5h. Cell lysates were assayed for luciferase activities and normalized for β-galactosidase activities. Statistical significance was calculated by independent t-test on N=3 experiments. Results are shown as mean ± SEM. ***P<0.001 vs VEH

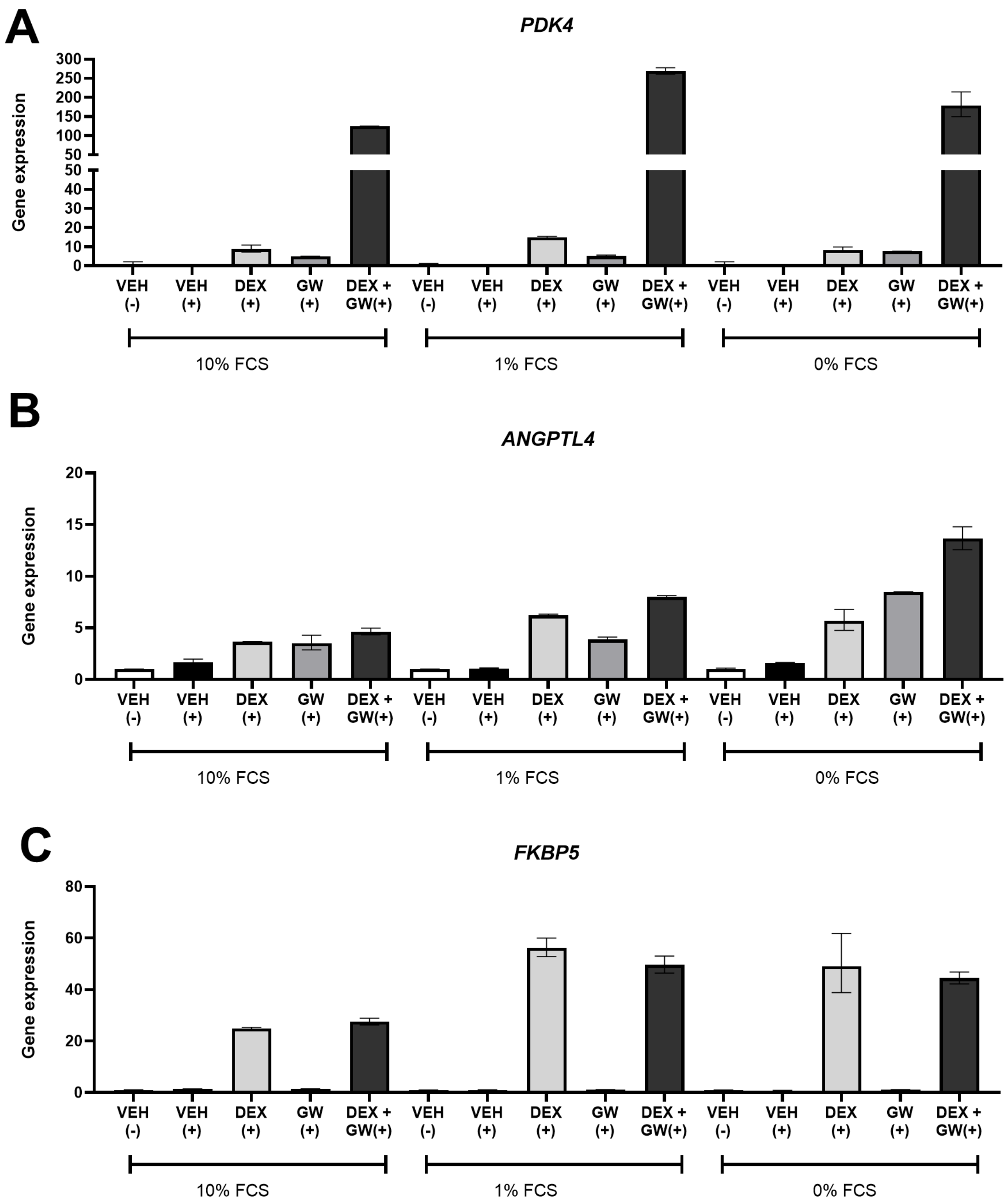


**Supplementary figure 2 - Serum starvation appears to induce stronger induction of shared GR-PPAR target genes in human fibroblast-like synoviocytes.** Human fibroblast-like synoviocytes were cultured in medium with 10%, 1% or 0% fetal calf serum (FCS) 16h before the experiment. Cells were pre-treated with (combinations of) dexamethasone (0.01µM, DEX), GW7647 (1µM, GW) or vehicle (VEH) for 1h, before stimulation with human TNFα (+) or medium (-) for an additional 23h. Expression of (shared) GR and PPAR target genes were analyzed by qPCR and shown as fold change relative to the TNF(-) group. No statistical significance was calculated (N=1 experiment). Results are shown as mean ± SD calculated on three technical replicates.

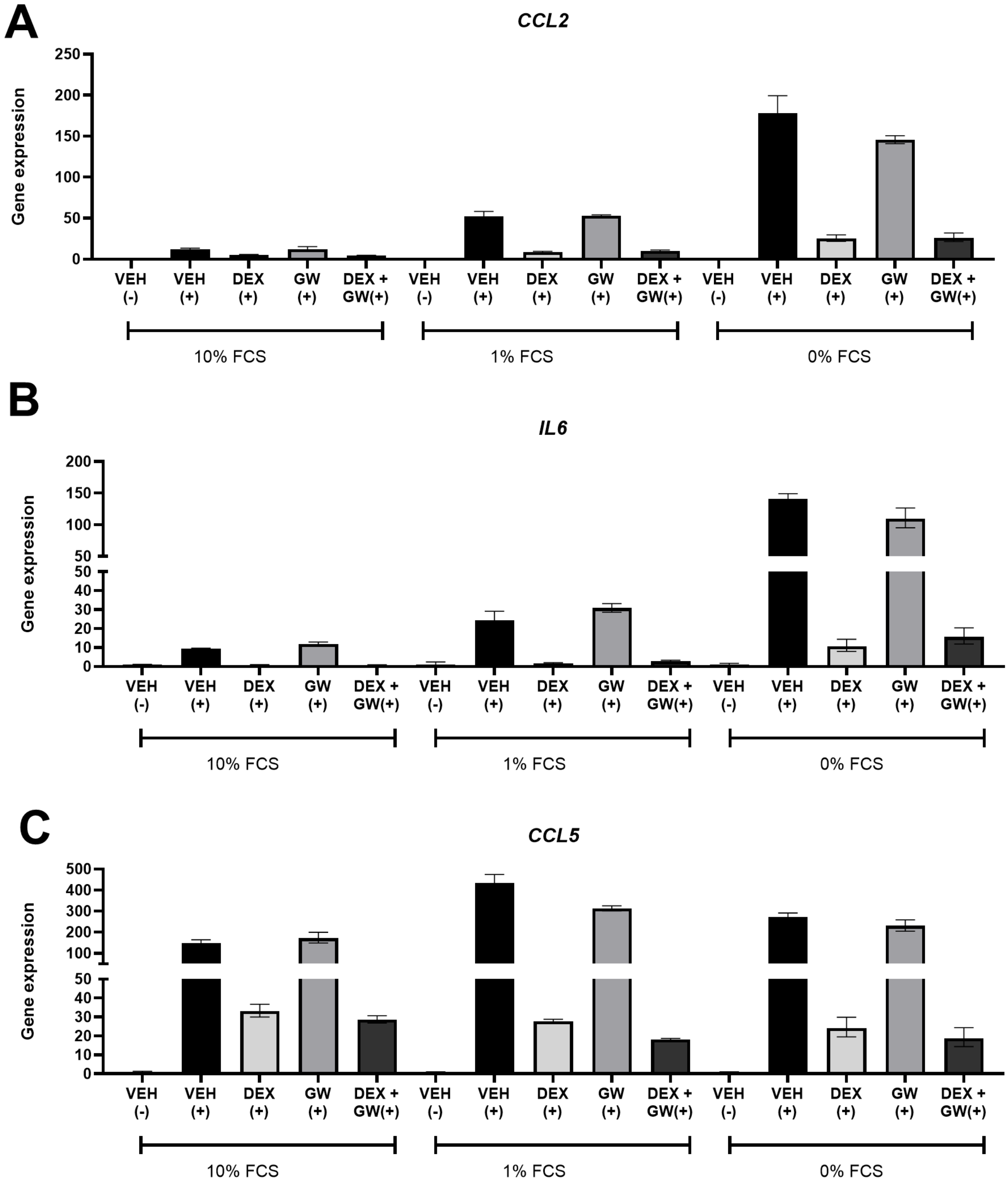


**Supplementary figure 3 - GW7647 does not differentially regulate pro-inflammatory gene expression in human fibroblast-like synoviocytes.** Human fibroblast-like synoviocytes from N=4 patients were cultured in medium with 10%, 1% or 0% fetal calf serum (FCS) 16h before the experiment. Cells were pre-treated with (combinations of) dexamethasone (0.01μM, DEX), GW7647 (1μM, GW) or vehicle (VEH) for 1h, before stimulation with human TNFα (+) or medium (-) for an additional 23h. Expression of (shared) pro-inflammatory genes were analyzed by qPCR and shown as fold change relative to the TNF(-) group. No statistical significance was calculated (N=1 experiment). Results are shown as mean ± SD calculated on three technical replicates.

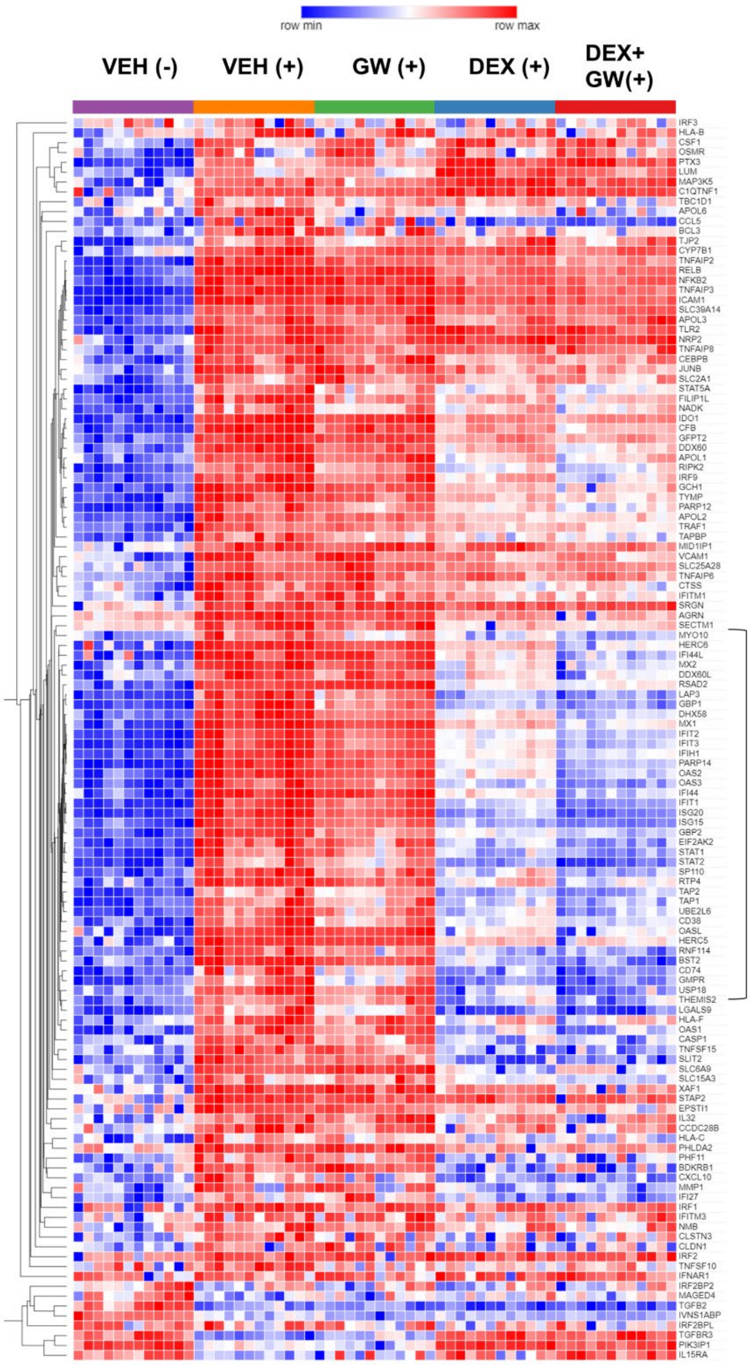


**Supplementary figure 4 - GW7647 enhances suppression of interferon-stimulated proteins after dexamethasone in human fibroblast-like synoviocytes.** Human fibroblast-like synoviocytes (FLS) from N=4 patients were serum-starved for 16h before the experiment. Cells were pre-treated with (combinations of) dexamethasone (0.01µM, DEX), GW7647 (2.5µM, GW) or vehicle (VEH) for 1h, before stimulation with human TNFα (+) or medium (-) for an additional 23h, followed by shotgun proteomics. All interferon-stimulated-proteins (ISPs) identified in the proteomics dataset were selected for heatmap visualization, and hierarchical clustering was exclusively applied on ISPs, using hierarchical clustering of z-scored, quantified log-transformed expression values.

**Supplementary Table 1: Primer sequences that were used for RT-qPCR analysis.**

| Species | Gene | Primer fw (5' to 3') | Primer rev (5' to 3') |
|---|---|---|---|
| *Mus musculus* | *Angptl4* | GGAAAGAGGCTTCCCAAGATG | CGTTGGGAGTCAAGCCAATG |
| *Mus musculus* | *B2m* | CATGGCTCGCTCGGTGAC | CAGTTCAGTATGTTCGGCTTCC |
| *Mus musculus* | *Ccl2* | CAAGAAGGAATGGGTCCAGA | AGACCTTAGGGCAGATGCAG |
| *Mus musculus* | *Ccl5* | TTTGCCTACCTCTCCCTCG | CGACTGCAAGATTGGAGCACT |
| *Mus musculus* | *Fkbp5* | TGAGGGCACCAGTAACAATGG | CAACATCCCTTTGTAGTGGACAT |
| *Mus musculus* | *Gapdh* | AACTTTGGCATTGTGGAAGG | ACACATTGGGGGTAGGAACA |
| *Mus musculus* | *Hprt1* | TGACACTGGCAAAACAATGCA | GGTCCTTTTCACCAGCAAGCT |
| *Mus musculus* | *Icam1* | TGCCTCTGAAGCTCGGATATAC | TCTGTCGAACTCCTCAGTCAC |
| *Mus musculus* | *Il1b* | CAGGCAGGCAGTATCACTCA | TGTCCTCATCCTGGAAGGTC |
| *Mus musculus* | *Il6* | GTTCTCTGGGAAATCGTGGA | CAGAATTGCCATTGCACAAC |
| *Mus musculus* | *Mmp3* | GCA GTT TGC TCA GCC TAT CC | GAG TGT CGG AGT CCA GCT TC |
| *Mus musculus* | *Nfkb1* | CGAGACTTTCGAGGAAATACCC | GTCTGCGTCAAGACTGCTACA |
| *Mus musculus* | *Nfkb2* | TGACTGTGGAGCTGAAGTGG | AAGGAGGCGAGTAAGAGTTGG |
| *Mus musculus* | *Pdk4* | CACCACATGCTCTTCGAACTCT | AAGGAAGGACGGTTTTCTTGATG |
| *Mus musculus* | *Tsc22d3* | GGCCCTAGACAACAAGATTGAG | CACGAATCTGCTCCTTTAGGAC |
| *Homo sapiens* | *CCL2* | CAGCCAGATGCAATCAATGCC | TGGAATCCTGAACCCACTTCT |
| *Homo sapiens* | *CCL5* | TGCCCACATCAAGGAGTATTT | TTTCGGGTGACAAAGACGA |
| *Homo sapiens* | *GAPDH* | TGCACCACCAACTGCTTAGC | CATGGACTGTGGTCATGAG |
| *Homo sapiens* | *HERC6* | CTGCCAAGCCTAAACCTGAG | CCAATGTCATCAGCAGCATC |
| *Homo sapiens* | *HERC6* | CTGCCAAGCCTAAACCTGAG | CCAATGTCATCAGCAGCATC |
| *Homo sapiens* | *HPRT1* | TGACACTGGCAAAACAATGCA | GGTCCTTTTCACCAGCAAGCT |
| *Homo sapiens* | *IFI44L* | TGCACTGAGGCAGATGCTGCG | TCATTGCGGCACACCAGTACAG |
| *Homo sapiens* | *IL6* | GACAGCCACTCACCTCTTCA | AGTGCCTCTTTGCTGCTTTC |
| *Homo sapiens* | *ISG15* | GGAATAACAAGGGCCGCAGCAG | AGGTCAGCCAGAACAGGTCGTC |
| *Homo sapiens* | *MX1* | GGTGGTCCCCAGTAATGTGG | CGTCAAGATTCCGATGGTCCT |
| *Homo sapiens* | *PDK4* | TTCTCAGGGGAACACCACCTCCTC | CGGGCAACAGTTGAACACCAGG |

#### SUPPLEMENTARY MATERIALS AND METHODS

##### Animal studies

*Arthritis severity scoring*

The clinical severity of arthritis per paw was graded daily according to standard evaluation procedures: 0, normal paws; 0.5, edema and erythema of only one interphalangeal joint; 1, edema and erythema of at least two interphalangeal joints or the metacarpophalangeal joint or the carpal/tarsal joint; 2, edema and erythema of two joint types; 3, edema and redness involving the entire paw; total clinical severity score being the sum of the clinical severity scores of all four paws.

*Serum anti-IgG1 and -IgG2 antibody analysis*

To evaluate serum anti-CII antibody levels, blood samples were collected at the end of the CIA experiments and were allowed to clot at room temperature. Upon centrifugation, they were stored at -20°C. At the day of the ELISA, 96-well half area microplates (Greiner Bio One) were coated with CII diluted in coating buffer (1.26 µl/ml) for 2 hours at 37°C. Blocking of nonspecific binding was done with 0.1 % casein in PBS without azide for 1 hour at 37°C. Samples were thawed and added to the wells in two dilutions (1/10 and 1/100 in blocking buffer). Upon overnight incubation at 4°C, detection antibodies coupled to HRP (IgG1-HRP and IgG2$\alpha$-HRP, Southern Biotech) were added for 1 hour at room temperature. Finally, TMB substrate solution (OptEIA TMB substrate Reagent set, BD biosciences) was added and the reaction was stopped by addition of 1 M H2SO4. OD values were measured at 450nm (Multiskan RC, Thermo Labsystems).

##### Proteomic profiling

*Preparation of LC-MS/MS samples*

FLS cells were seeded in three technical replicates in 12-well plates (100,000 cells/well) and, 24h later, were serum deprived for 16h prior to compound stimulations. FLS cells were washed with ice-cold PBS, collected with 5% SDS / 50 mM triethylammonium bicarbonate (TEAB), snap-frozen and stored at -80 °C until further processing. The lysate was transferred to a 96-well PIXUL plate and sonicated with a PIXUL Multisample sonicator (Active Motif) for 5 minutes with default settings (Pulse 50 cycles, PRF 1 kHz, Burst Rate 20 Hz). Next, samples were centrifuged for 15 minutes at 2,204 xg at RT to remove insoluble components. Proteins were reduced and alkylated by addition of 10 mM Tris(2-carboxyethyl)phosphine hydrochloride and 40 mM chloroacetamide and incubation for 10 minutes at 95°C in the dark. Phosphoric acid was added to a final concentration of 2.75% and subsequently samples were diluted 7-fold with binding buffer containing 90% methanol in 100 mM TEAB, pH 7.55 (binding buffer). After loading the samples to S-trap micro columns (Protifi) using centrifugation for 1 min at 1,500 xg, the columns were washed 3 times with 150 µl binding buffer using the same centrifugation settings. Digestion of the proteins was done by addition of 20 µl 50mM TEAB containing 1 µg of trypsin and incubation overnight at 37°C. Peptides were eluted in 3 times, first with 40 µl 50 mM TEAB, then with 40 µl 0.2% formic acid (FA) in water and finally with 40 µl 0.2% FA in water/acetonitrile (ACN) (50/50, v/v). Eluted peptides were dried completely by vacuum centrifugation. The dried peptides were redissolved in 20 µl 0.1% trifluoroacetic acid in water/acetonitrile (98/2, v/v), diluted 100-fold with 0.1% formic acid (FA) of which 20 µl was loaded on Evotips (Evosep, P/N EV2011, Odense, Denmark) according to the manufacturer's instructions. All loaded Evotips were stored in 0.1% FA at 4°C until LC-MS/MS analysis could be started.

*LC-MS/MS analysis*

FLS cells were washed with ice-cold PBS, collected with 5% SDS / 50 mM triethylammonium bicarbonate (TEAB), snap-frozen and stored at -80 °C until further processing. Samples were run in data-independent parallel accumulation serial fragmentation (DIA-PASEF) mode on an Evosep One LC-system (Evosep) in-line connected to a timsTOF SCP (Bruker, Belgium). Peptides were analyzed with the 20 SPD whisper method using the Aurora Gen3 Elite column (15cm x 75 µm I.D., 1.7µm beads, Evosep), heated to 50°C. Peptides were eluted from the column through the predefined 20SPD whisper gradient consisting of 0.1% FA in LC-MS-grade water as solvent A and 0.1% FA in ACN as solvent B. Eluting peptides were measured in positive polarity with a full-scan range of 100 m/z to 1700 m/z. The trapped ion mobility spectrometry (TIMS) was operated at a fixed duty cycle close to 100%, a ramp and accumulation time of 100 ms, ranging from 1/K0 =0.64 Vscm2 to 1/K0 =1.50 Vscm2. Collision energy was linearly ramped as a function of the inverse mobility from 20 eV at 1/K0=0.60Vscm2 to 59 eV at 1/K0 = 1.60 Vscm2. A DIA-PASEF mass range of 400 Da to 1000 Da was used in a mobility range of 1/K0 =0.64 Vscm2 to 1/K0 =1.37 Vscm2 using a window size of 25 Da according to supplementary table S1, resulting in a cycle time of 0.96s.

*LC-MS/MS data analysis*
LC-MS/MS runs of all samples were searched together using the DiaNN algorithm (version 1.8.1). Spectra were searched against the human protein sequences in the Swiss-Prot database (database release version of 2023_08), containing 20,423 sequences. Enzyme specificity was set as C-terminal to arginine and lysine, also allowing cleavage at proline bonds with a maximum of 2 missed cleavages. Variable modifications were set to oxidation of methionine residues and acetylation of protein N-termini. Fixed modification was set to carbamidomethylation of cysteine residues. Default settings were mainly used, except for the addition of a 400-1000 m/z precursor mass range filter and MS1 and MS2 mass tolerance was set to 15 and 20 ppm, respectively. Further data analysis of the shotgun results was performed with an in-house script in the R programming language (version 4.2.2). Protein expression matrices were prepared as follows: the DIA-NN main report output table was filtered at a precursor and protein library q-value cut-off of 1% and only proteins identified by at least one proteotypic peptide were retained. After pivoting into a wide format, iBAQ intensity columns were then added to the matrix using the DIAgui's R package get_IBAQ function (26). Groups of 7,650 proteins were reliably quantified using protein group maximum likelihood quantification (PG.MaxLFQ), with at least 3 valid values in one of the experimental conditions.

Proteomics data were log2-normalized and missing protein intensity values were imputed by randomly sampling from a normal distribution centered around each sample's noise level (package DEP) (27). Principle component analysis was performed using R. To compare protein abundance between pairs of sample groups, statistical testing for differences between 2 group means was performed, using the package limma (28), with a false discovery rate (FDR) < 0.05, log fold change of ≥ 2- or ≤ 0.5-fold (|log2FC| ≥ 1) and adjusted p-value < 0.05. Ingenuity pathway analysis (IPA) was performed for gene set enrichment analysis and network analysis on all proteins with an FDR and adjusted *p*-value < 0.05, without applying an additional log-fold change threshold ($|\log_2 FC| \geq 1$). Heatmap visualization, and hierarchical clustering was exclusively applied on all interferon-stimulated genes-proteins present in the dataset, using hierarchical clustering of z-scored log2 PG.MaxLFQ intensities (metric: one minus pearson correlation; linkage method: average; cluster: columns).

**Western Blotting and supernatant analysis**
FLS cells were seeded in 6-well plates (300,000cells/well, respectively) and 24h later, serum deprived for 16h prior to compound stimulations. Supernatants were collected and diluted 25x for measurement of CCL2 levels using the MCP-1/CCL2 Uncoated ELISA kit (Invitrogen). Total cell lysates were prepared using 1x SDS sample buffer (50mM Tromethamine (Tris) pH 6.8; 2% SDS; 10% glycerol; bromophenol blue and 100mM DL-Dithiothreitol;, freshly added).

Samples were incubated at 95◦C for 5min and separated on a SDS-PAGE gel and subsequently blotted onto a nitrocellulose membrane (Whatman, Dassel, Germany). Immunoblotting was performed according to the standard protocol of Santa Cruz (Santa Cruz, CA, USA). Primary antibodies used were targeting GR (1/2000, sc-393232, Santa-Cruz), PPARα (1/2000, sc-398394, Santa-Cruz) and α-tubulin (1/4000, Sigma-Aldrich). As secondary antibodies, we used species-specific HRP-conjugated antibodies (cat nr: NA931, NA934, GE-Healthcare) after which ECL prime Western Blot Detection Reagent (Sigma-Aldrich) was added to visualize protein signal via Western Lightning (PerkinElmer, Waltham, MA, USA). To quantify bands obtained via Western analysis, background intensity was subtracted from the mean intensity per lane using ImageJ Fiji [4]. To correct for in-between blot differences in intensities, fold change was calculated per blot, using the VEH+ group as reference.

**RT-qPCR**

For L929sA experiments, cells were seeded in 24-well plates (30,000cells/well) and 24h later serum-deprived; in case of the IFN experiments, cells were additionally treated with IFNAR antibody (6.5ug/mL, MAR1-5A3, Leinco), for 16h prior to compound treatments. Total RNA was isolated using the Qiashredder and Rneasy mini kits (Qiagen). Reverse transcription (RT) was performed using the Biotech Rabbit cDNA synthesis kit. For FLS experiments, cells were seeded in 24-well plates (50,000cells/well) and 24h later, serum-deprived for 16h prior to compound stimulations. Total RNA was isolated using the Qiashredder and Rneasy micro kits (Qiagen). Reverse transcription (RT) was performed using the iScript cDNA synthesis kit (Bio-Rad). For mouse experiments, knee synovia were dissected and stored in RNA Later (Qiagen) until the day of RNA isolation by the RNeasy mini kit (Qiagen). cDNA was synthesized using the QuantiTect Reverse Transcription kit (Qiagen), according to manufacturer's instructions.
For all experiments, RT-qPCR was performed using SYBR-Green I Master Mix (Roche) and the LightCycler480 (Roche Diagnostics). Primers were tested for high efficiency (90%-110%) and for amplification of a single PCR product. All primer sequences are listed in **Supplementary Table S1**, and fold change expression was calculated using the $2^{-\Delta\Delta CT}$ method using *Gapdh, Hprt1* and *B2M* as housekeeping genes for L929sA cells and murine synovia, and *HPRT1* and *GAPDH* for FLS.
